# A weighted and cumulative point system for accurate scoring of intestinal pathology in a piglet model of necrotizing enterocolitis

**DOI:** 10.1101/2024.01.05.574327

**Authors:** Simone Margaard Offersen, Nicole Lind Henriksen, Anders Brunse

## Abstract

**Background:** Necrotizing enterocolitis (NEC) is a serious condition, primarily affecting premature infants, in which a portion of the gut undergoes inflammation and necrosis. Symptoms of NEC are unspecific, and together with a rapid progression, the disease remains a significant concern. The preterm pig develops NEC spontaneously, making it a suitable model for exploring novel treatments. During piglet necropsy, NEC-lesions closely resemble the pathologies found during surgery or autopsy of preterm infants. As such, the systematic gross inspection enables direct evaluation of gut lesions, which is not possible in the medical preterm patient. Here, we introduce a revised intestinal scoring system with an expanded score range and more detailed descriptive features to accurately describe the diversity of NEC-lesions in the preterm piglet model.

**Methods:** We included 333 preterm piglets from four separate experiments, each delivered via cesarian section at 90% gestation. The pigs were fed either a gently processed (GP) or harshly processed (HP) milk formula for 96 hours and were subsequently euthanized. At necropsy, the gastrointestinal tract was assessed with 1) an established 6-grade scoring system and 2) a systematic and descriptive approach focusing on the distribution and severity of hyperemia, hemorrhage, pneumatosis intestinalis (intramural gas), and necrosis. Lesion biopsies were sampled for cytokine measurement and a subset (*n* = 62) was sampled for histopathological assessment.

**Results:** The systematic and descriptive registrations were evaluated and converted into a weighted and cumulative point (WCP) score. Compared to the 6-grade score, the WCP score enabled a higher discrepancy in severity levels, especially among organs with more prominent NEC lesions. IL-1β in small intestinal lesions and both IL-8 and IL-1β in colon lesions correlated positively with the WCP scale. A histopathological grade system (0-8) was established and revealed mucosal lesions not recognized macroscopically. Finally, the WCP score showed a higher NEC-promoting effect of the HP formula compared to the GP formula.

**Conclusion:** The validation of the weighted and cumulative scoring system demonstrated an expanded score range, enhancing the accuracy in describing NEC-lesions in gastrointestinal segments of preterm pigs. This approach may increase the efficiency of preclinical NEC experiments.

## Background

Necrotizing enterocolitis (NEC) is a life-threatening gastrointestinal disease almost exclusively affecting preterm infants. The immature immune and digestive systems render preterm infants vulnerable to NEC, with factors such as formula feeding and bacterial colonization playing pivotal roles [1, 2]. On average, 5-10% of preterm neonates develop NEC with higher incidences among infants born very preterm (< 28 weeks of gestation) or with a very low birth weight (VLBW, < 1500 g) [3, 4]. The severe and rapid progression of NEC makes the disease a particular challenge for clinicians in the neonatal intensive care unit.

NEC is inherently characterized by inflammation and necrosis of the intestines. However, since the intestines are not easily assessed in the preterm human infant, early diagnosis is based on clinical and radiological findings rather than specific intestinal lesions. The clinical signs of NEC include abdominal distention and feeding intolerance followed by bloody stools, discolored abdominal wall, and respiratory distress. Radiological findings include portal venous gas and intestinal intramural gas (pneumatosis intestinalis), which is a pathognomonic feature of NEC [5, 6]. When signs of NEC are recognized, medical treatment is started with e.g. gastric decompression, bowel rest, and antibiotic treatment. Surgery is indicated if the response to medical therapy fails or if the clinical symptoms deteriorate [5, 7]. Only upon inspection of surgically resected or autopsied intestines, the actual presence of inflammation and necrosis can be assessed. The most frequent pathological sites of injury involve the ileum and colon and the observed lesions include dilated, hemorrhagic, or necrotic intestines, often accompanied by pneumatosis intestinalis [6].

The newborn pig is considered a suitable model for human infants in terms of gut development and disease since the anatomy and physiology of the piglet’s gastrointestinal tract closely resemble that of human infants [8–10]. When delivered prematurely at 90% gestation, piglets have a similar body size as preterm infants. Moreover, intestinal development, immune response, and susceptibility to NEC are largely comparable between the two species in a preterm state. Importantly, preterm piglets require no artificial stressors but develop variable degrees of NEC-like intestinal pathology spontaneously, with higher severity if fed infant formula. Moreover, the spectrum of clinical observations and gut pathology observed in autopsied preterm piglets closely resemble typical clinical and pathological findings in preterm infants diagnosed with NEC from feeding intolerance, abdominal distention, and intestinal hemorrhage to pneumatosis intestinalis, transmural necrosis, portal venous gas, and pneumoperitoneum in the most severe cases [8].

NEC-like lesions in preterm pigs are usually scored with a simple 6-grade system ranging from no lesions or hyperemia (score 1-2) to local or extensive hemorrhage (score 3-4) and, finally, to local or extensive necrosis and/or pneumatosis intestinalis (score 5-6) [8, 11]. While this approach is simple and easy to interpret, it combines diverse pathologies into a single categorical score and cannot describe lesions that are neither focal nor diffuse, potentially leading to a loss of important information and misclassification. Given that a comprehensive and accurate evaluation of intestinal lesions is a critical outcome measure in experimental NEC studies, we advocate for the adoption of a revised system with an expanded score range and more detailed descriptive features.

Here, we introduce a weighted and cumulative point (WCP) system to score intestinal lesions in pigs based on the registered extent of four separate pathologies: hyperemia, hemorrhage, pneumatosis intestinalis, and necrosis. We demonstrate how it correlates to levels of intestinal pro-inflammatory cytokines and a newly developed histopathological grading system. Lastly, we demonstrate the use with a data example comparing the NEC-promoting effect of two milk formulas.

## Methods

### *In vivo* experimental procedures and monitoring

Experimental procedures conducted during the animal experiments were approved by the Danish Animal Experiments Inspectorate (license number, 2020-15-0201-0052), under the guidelines from Directive 2010/63/EU of the European Parliament. Animal registrations originated from four separate studies conducted over two years. A total of 333 crossbred piglets ((Landrace × Yorkshire) × Duroc) were delivered by cesarean section from 15 sows at 90% gestation (106 days). After delivery, the piglets were stabilized and equipped with orogastric tubes for enteral feeding and umbilical arterial catheters for parenteral nutritional supplementation. Sex and birth weights were registered and the piglets were individually housed in preheated incubators as described previously [8].

All pigs received sow plasma by arterial infusion (16 mL/kg) on the first day of life instead of the normal colostrum immunization. Hereafter, the pigs were provided with declining levels of intraarterial parenteral nutrition (PN) during the study period (2-4 mL/kg^0.75^/d, Kabiven, Vamin, Fresenius-Kabi; Bad Homburg, Germany). Every third hour, the pigs were fed enteral boluses of milk formula. 263 piglets (12 litters) received a gently processed (GP) home-mixed formula composed of maltodextrin (50 g/L, Fantomalt; Nutricia, Denmark), caseinate (15 g/L, Miprodan 40; Arla Foods AMBA, Denmark), whey protein isolate (60 g/L, Lacprodan DI-9224; Arla Foods Ingredients, Denmark), medium- and long-chain triglycerides (75 g/L Liquigen and 20 g/L Calogen; Nutricia, Denmark), and vitamins (2 g/L, Phlexy Vits; Nutricia, Denmark). The remaining 70 piglets (3 litters) received a harshly processed (HP) liquid ready-to-drink infant formula (PreNAN Preemie; Nestlé, Switzerland) supplemented with extra whey protein (21 g/L, Lacprodan DI-9224; Arla Foods Ingredients, Denmark) and medium-chain triglycerides (21 g/L, Liquigen; Nutricia, Denmark). All four studies had pigs receiving GP formula and two of the studies had pigs receiving HP formula (Table 1). The formula volumes increased gradually from 24 mL/kg^0.75^/d on day 1 to reach 80-96 mL/kg^0.75^/d on day 4-5 (further specified in Supplementary Figure S1). The formula energy compositions can be found in Supplementary Table S1.

**Table 1:** Overview of sample size across studies, milk formulas, and outcomes.

|  | Study 1 | Study 2 | Study 3 | Study 4 | Total |
| --- | --- | --- | --- | --- | --- |
| *Formulas:* |  |  |  |  |  |
| GP | 84 | 64 | 44 | 71 | 263 |
| HP | 0 | 17 | 0 | 53 | 70 |
| *Outcomes:* |  |  |  |  |  |
| Macroscopic score | 84 | 81 | 44 | 124 | 333 |
| Lesion cytokines | 82 | 81 | 0 | 118 | 281 |
| Histology grade | 62 | 0 | 0 | 0 | 62 |

GP = gently processed milk formula, HP = harshly processed milk formula.

The pigs were kept under close monitoring by experienced personnel at least every third hour. Pigs that presented with clinical signs of NEC (e.g. abdominal distension, severe rectal bleeding, or respiratory distress) were immediately euthanized. On day five, all pigs were deeply anesthetized and subsequently euthanized with a lethal dose of pentobarbital.

### Post-mortem macroscopic lesion scoring and tissue collection

Following euthanasia, the gastrointestinal (GI) tract was extracted and released from the mesentery. The stomach and colon were separated from the small intestine at the pyloric orifice and the ileocecal junction, respectively. The stomach and small intestine were emptied for digesta and the colon was isolated from the rectum. Hereafter, each of the three GI segments was evaluated for macroscopic lesions with two different methods: 1) an established categorical 6-grade system and 2) a systematic, descriptive approach focusing on lesion distribution and pathological severity (i.e. a spectrum from hyperemia to necrosis).

The 6-grade system has been used extensively in preterm piglet studies and was scored as follows: 1 = normal appearance; 2 = hyperemia; 3 = local hemorrhage; 4 = extensive hemorrhage; 5 = local necrosis and/or pneumatosis intestinalis, and 6 = extensive necrosis and/or pneumatosis intestinalis [8, 11, 12]. For the systematic, descriptive method, the presence of hyperemia, hemorrhage, pneumatosis intestinalis, and necrosis were registered separately. Each pathology type was described by one of four ordinal levels, based on the spatial extent of the lesion: none, mild, moderate, or severe. Intestinal perforation was defined as severe necrosis. Pictures were captured from all intestinal regions.

### Cytokine measurement and histopathological grading of intestinal lesions

The worst macroscopic lesions from the small intestine and colon were snap-frozen or immersed in 4% paraformaldehyde for later cytokine analysis and histopathological assessment, respectively. If the lesion extent was very local, cytokine analysis was prioritized. If no lesions were observed, the tissue was sampled from a random location. Interleukin (IL) 8, IL-1β, and tumor necrosis factor (TNF) α levels were analyzed in tissue homogenates from Study 1, 2, and 4 by commercial porcine ELISA kits (R&D Systems, Abingdon, Oxfordshire, United Kingdom) according to the manufacturer’s instructions and expressed as picograms per milligram of wet tissue [13]. Formalin-fixed small intestine and colon lesion biopsies from Study 1 were embedded in paraffin and two serial 3-5 μm sections were made and stained with hematoxylin and eosin. The sections were assessed by an experienced investigator blinded to the investigation. Sample sizes are presented in Table 1.

### Statistical analysis

Statistical analyses were performed in R version 4.2.2. All correlations were analyzed with Spearman’s rank correlation coefficient. A linear model was used to analyze the effect of pathology type on cytokine levels. Here, dummy variables (0/1) of each pathology type (hyperemia, hemorrhage, pneumatosis intestinalis, and necrosis) were constructed and included as covariates together with the study origin. Cytokine levels were log10-transformed, and the main effect of each type was analyzed with an ANOVA test. WCP scores were analyzed with a proportional ordered logistic regression model (polr function in MASS package) with formula, birth weight, and study origin (1-4) as covariates. The main effect of formula was analyzed using the likelihood ratio test. The effect of sex was analyzed with an identical approach. The effect of milk formula on lesion cytokine levels was analyzed with a Mann-Whitney U test. Incidences of lesion types were compared with Fisher’s extract test. *P*-values below 0.05 were considered statistically significant.

## Results

### Development of a weighed and cumulative point system

After completion of all studies, the systematic descriptive records were assessed across GI segments of all animals. Hyperemia was predominantly observed in the small intestines, while hemorrhage was more frequently noted in stomachs and colons (Supplementary Fig. S2A). This was particularly pronounced in colons, where 59% had some degree of bleeding. Approximately 30% of stomachs and colons had observations of pneumatosis or necrosis. Importantly, 30% of stomachs, 11% of small intestines, and 44% of colons had more than one pathology type registered (Supplementary Fig. S2B). This prompted the development of a scoring system that 1) integrates all observations and 2) differentiates between mild and severe pathologies. Therefore, a weighted point system was developed with increasing values based on the severity of both the type and extent of the lesions (Table 2). The weighting of pathology type was based on the established categorical 6-grade score that ranges severities from hyperemia to hemorrhage to pneumatosis intestinalis, and necrosis [8, 11]. The lesion extent was expanded to three levels (mild, moderate, and severe) instead of the two levels in the 6-grade score (local and extensive). All degrees of hyperemia were weighted equally due to the limited correlation with NEC pathology observed in infants.

**Table 2:**
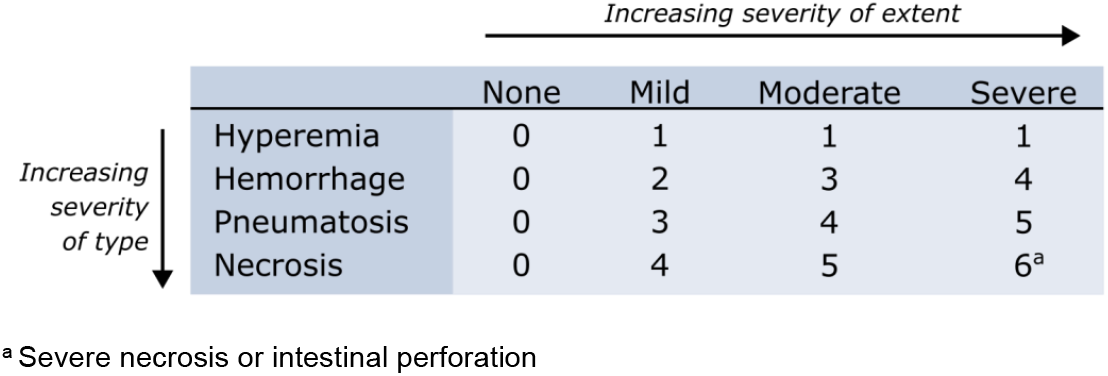
Weighted points based on severity of four pathology types and four lesion extent levels.

The cumulative score per GI segment was calculated as the sum of the weighted points for hyperemia, hemorrhage, pneumatosis intestinalis, and necrosis. Simultaneous registration of hyperemia and severe hemorrhage did not occur, resulting in a maximum allocation of 15 points per organ. Examples of weighted and cumulative point (WCP) scores from stomachs, small intestines, and colons are presented in Fig. 1, 2, and 3, respectively.

**Figure 1.**
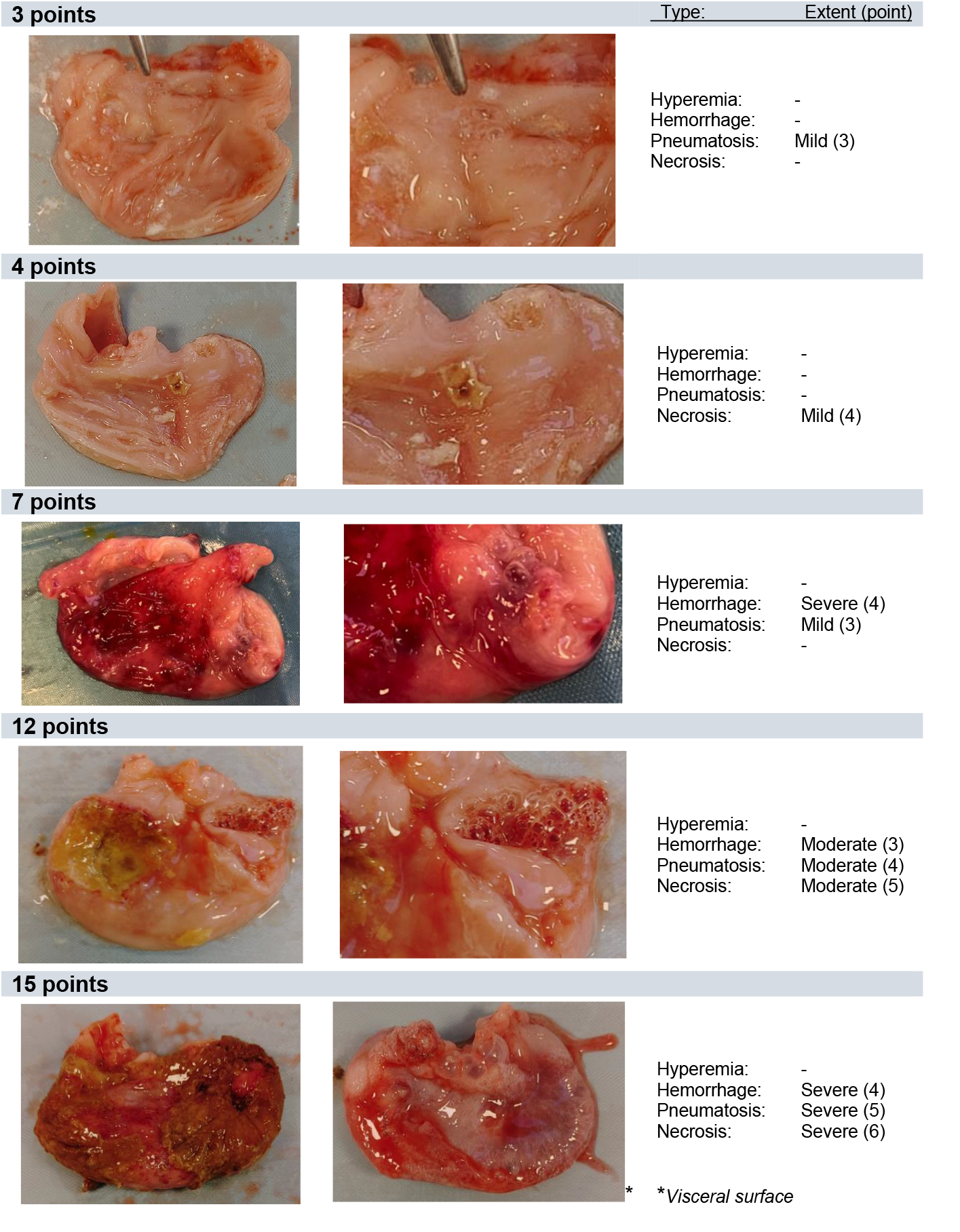
Weighted and cumulative score examples of the stomach. Unless otherwise stated, the organ is seen from the luminal surface.

**Figure 2.**
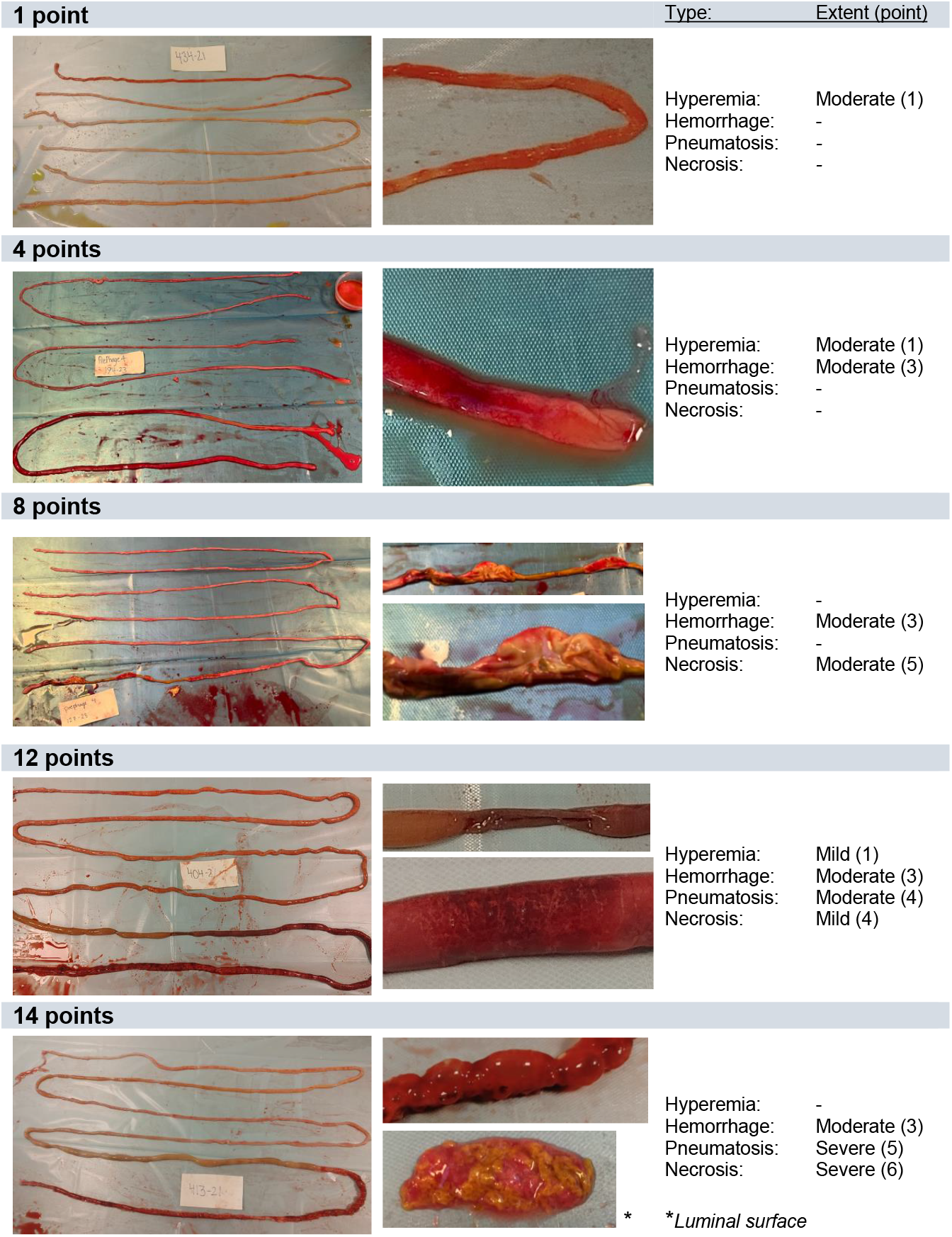
Weighted and cumulative score examples of the small intestine. Unless otherwise stated, the tissue is seen from the visceral surface. On left-sided pictures, the intestines are displayed with the proximal part above and distal part below.

**Figure 3.**
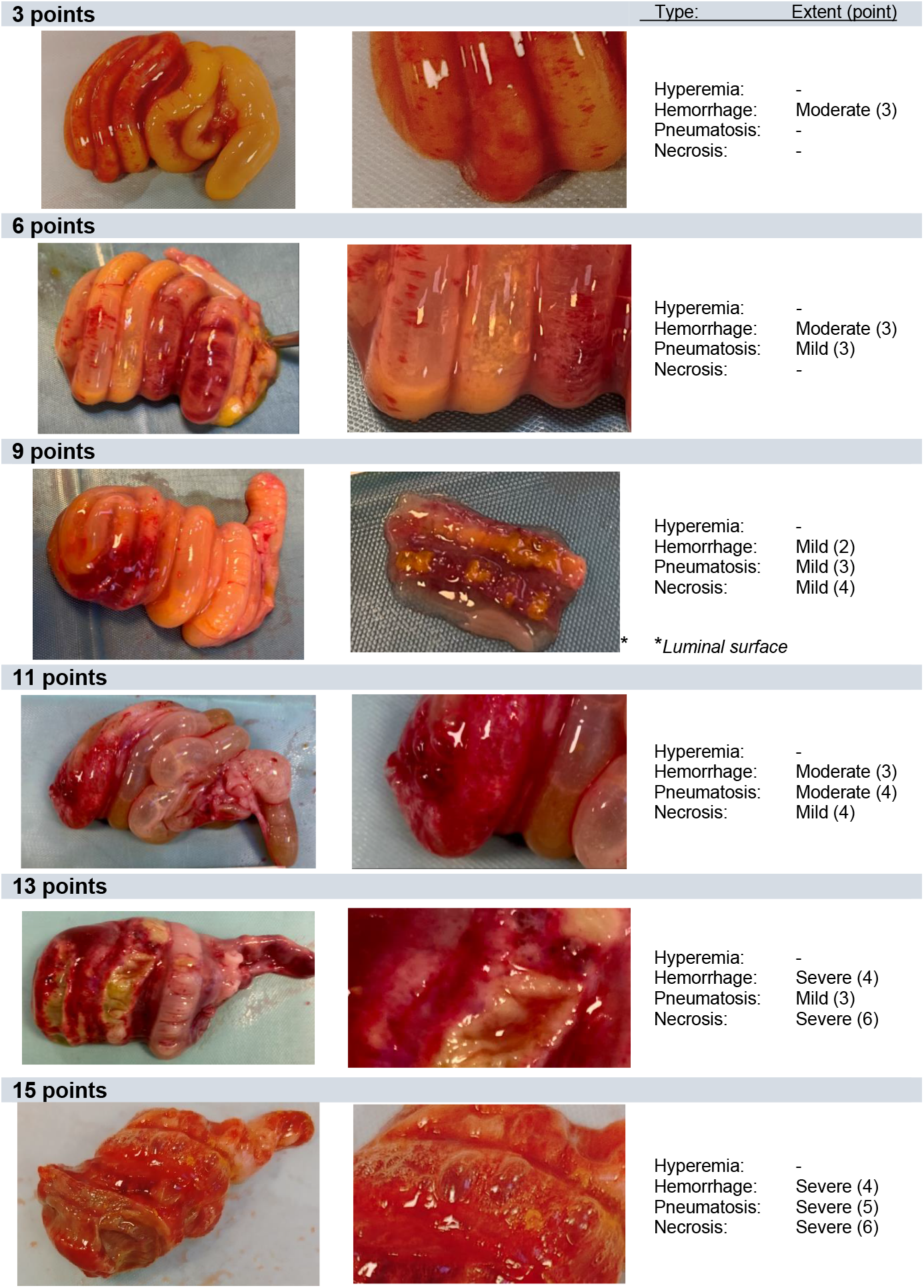
Weighted and cumulative score examples of the colon. Unless otherwise stated, the tissue is seen from the visceral surface.

### The weighted and cumulative point score allows higher discrepancy of NEC lesions

The macroscopic scoring was analyzed using both the categorical 6-grade score and the newly developed WCP score. When comparing the two, we observed that the WCP score permits a greater variation, particularly within the categorical grade 5-6 range (Fig. 4). Thus, the WCP score adds a discrepancy in severity levels among organs harboring more pronounced NEC-like lesions.

**Figure 4.**
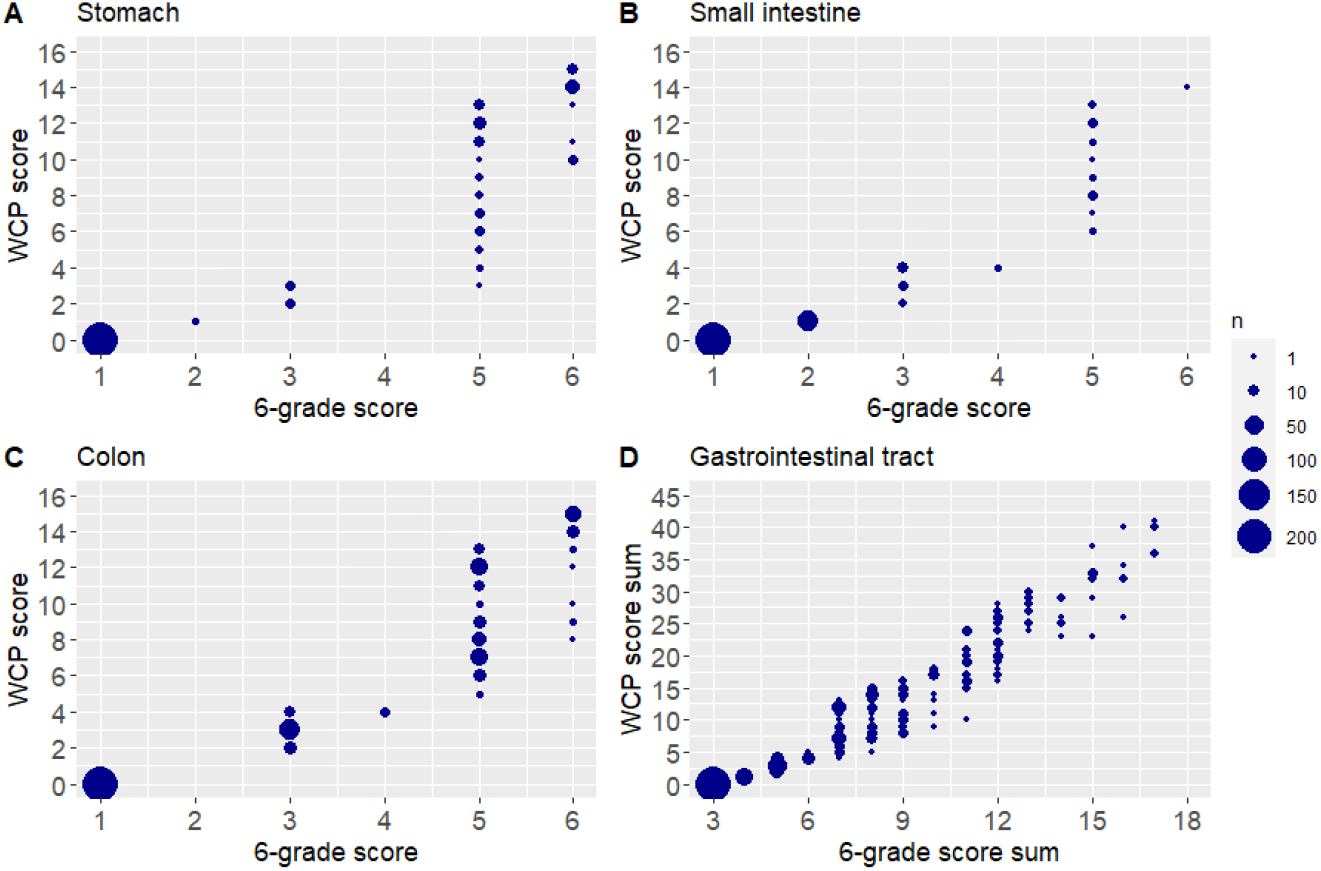
Correlation between the 6-grade system and weighted and cumulative point (WCP) score in each of the three gastrointestinal (GI) sections separately; the stomach (A), small intestine (B), and colon (C), as well as the sum across the gastrointestinal tract (D). *n* = 333 pigs.

### High-range scores correlate to elevated pro-inflammatory cytokines

During the inspection of small intestines and colons, tissue biopsies were collected from the most severely affected regions for measurement of three pro-inflammatory cytokines. TNF-α levels were below the detection limit in 77% of small intestinal biopsies and 92% of colon biopsies and were therefore excluded from the analysis. In small intestines, higher WCP scores correlated positively to IL-1β, whereas IL-8 showed equally high levels in both normal and affected tissues (Fig. 5A-B). Analyzing each pathology type separately revealed that the macroscopic presence of necrosis significantly increased both IL-8 and IL-1β levels (Supplementary Table S2). Notably, there was a 4.6-fold increase in IL-1β levels when necrosis was present. Hemorrhage increased IL-1β by 67% but did not affect IL-8. In contrast, hyperemia and pneumatosis intestinalis *reduced* IL-8 and did not affect IL-1β. Consequently, interpreting changes in small intestinal IL-8 levels should be approached with caution, as different pathology types demonstrate counteracting effects.

**Figure 5.**
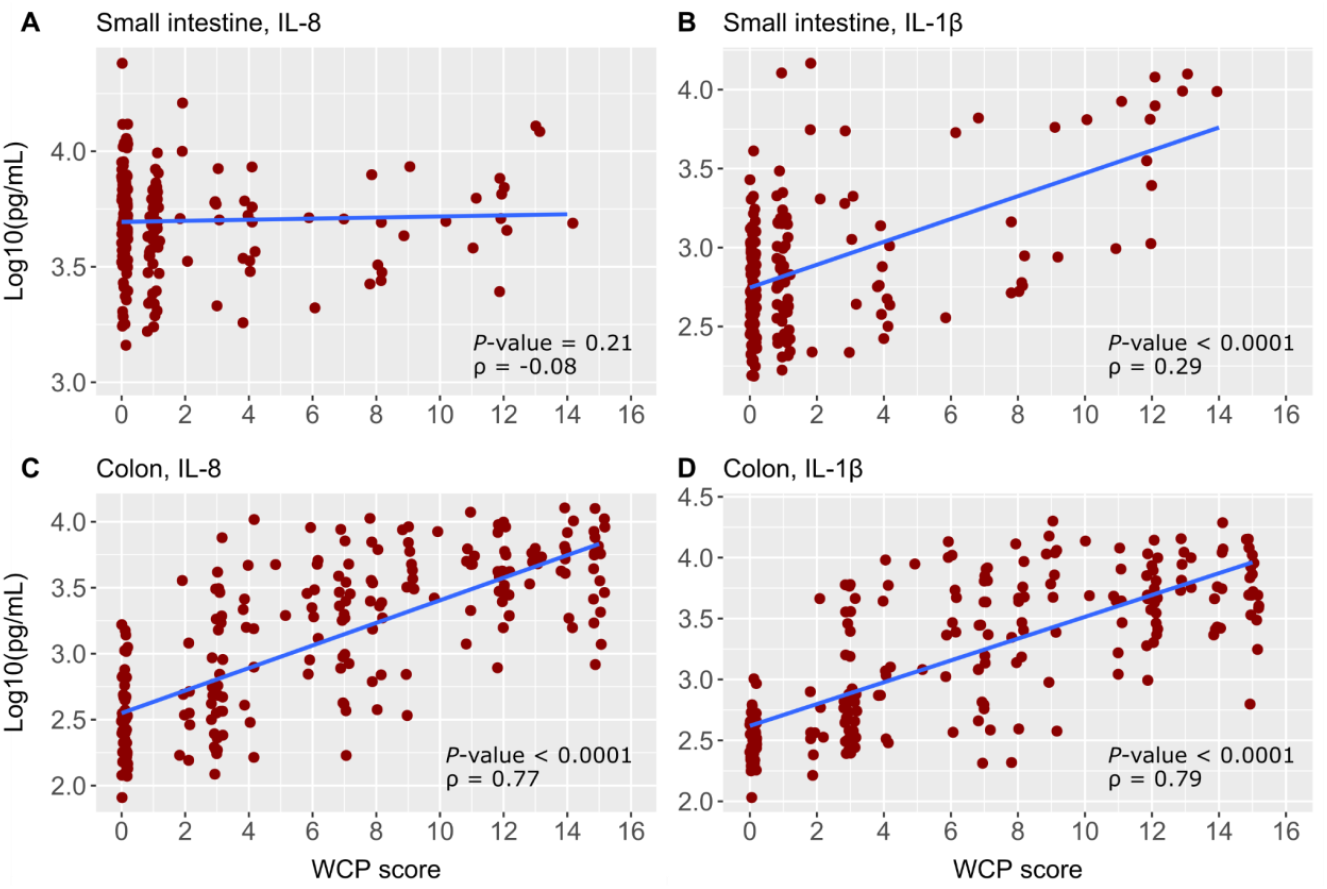
Correlation between weighted and cumulative point (WCP) scores and cytokine measurements using Spearman’s rho (ρ). A-B. Correlation between WCP scores in the small intestine and interleukin (IL)-8 levels (A) or IL-1β levels in small intestinal lesions (B). C-D. Correlation between WCP score in colons and IL-8 levels (C) or IL-1β levels (D) in colon lesions (*n* = 281).

For colons, both IL-8 and IL-1β displayed ascending trends with increased WCP scores (Fig. 5C-D). Since local cytokine measurements cannot capture overall organ involvement, they cannot differentiate between high-range scores of 10-15. When examining the presence of each pathology type, hemorrhage, pneumatosis intestinalis, and necrosis all significantly increased the level of both IL-8 and IL-1β in colons (Supplementary Table S2). Cytokine levels increased 2-2.7-fold with the registration of each of these types. There was no effect of hyperemia. Collectively, IL-1β in small intestinal lesions and both IL-8 and IL-1β in colon lesions can serve as paraclinical supportive data for an NEC assessment at the local tissue level. The good correlation between elevated WCP scores and cytokine levels verifies the WCP scale.

### Lesion histology supports macroscopic inspection

Histological sections of small intestinal and colonic lesions from *Study 1* were assessed and all types of observations were registered across all animals. Subsequently, these observations were categorized into severity grades ranging from 0 (normal appearance) to 8 (transmural ulceration). Given the distinct microscopic structures of the small intestinal and colonic mucosae, the lesions exhibited varied morphologies based on their location. Consequently, the definitions of grades 1-7 differed slightly between the organs. The criteria for small intestinal and colonic grading are listed in Table 3. Picture examples of each grade are presented in Fig. 6.

**Table 3.** Histopathological grading criteria.

| Grade | Small intestine | Colon |
| --- | --- | --- |
| 0 | Normal appearance | Normal appearance |
| 1 | Submucosal congestion | Exhausted goblet cells |
| 2 | Submucosal and villus congestion | Submucosal hemorrhage with or without cell infiltration |
| 3 | Local lifting or disruption of villus tip epithelium | Lifting of epithelial layer or local disruption of epithelium |
| 4 | Pronounced lifting or disruption of epithelium | Mucosal pneumatosis intestinalis without ulceration |
| 5 | Villus stunting | Local mucosal ulceration |
| 6 | Local mucosal ulceration | Pronounced mucosal ulceration |
| 7 | Pronounced mucosal ulceration | Widespread mucosal ulceration |
| 8 | Transmural ulceration | Transmural ulceration |

**Figure 6.**
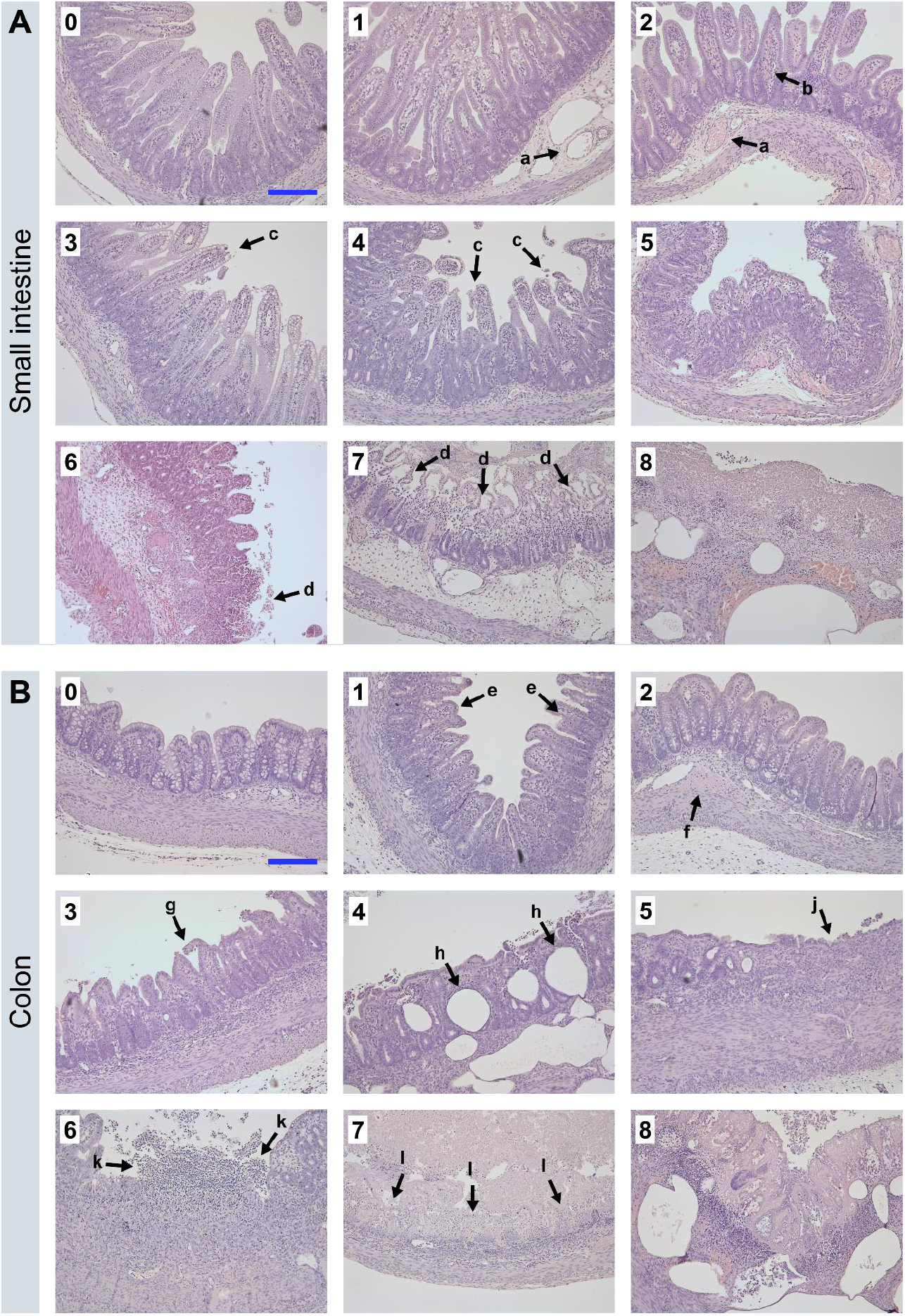
Histopathological grading of small intestines and colons. A) Histological changes in the small intestine with submucosa (a) and villus (b) congestion, disruption of the villus epithelium (c), and mucosal ulceration (d). B) Histological changes in the colon with exhausted goblet cells (e), submucosal hemorrhage (f), disruption of colon epithelium (g), mucosal pneumatosis intestinalis (h), and mucosal ulceration (i). Scale bar = 50 μm.

When examining the distribution of the macroscopic WCP scores, most stomachs and small intestines had no lesions (63% and 68%, respectively), resulting in a median score of 0 for both sections (Fig. 7A). The remaining WCP scores tended to be higher in the stomachs and lower in the small intestines. The colons displayed a higher level of macroscopic pathology, with only 41% being free of lesions and a median score of 3 points. The remaining colons were distributed relatively evenly on the point scale with the highest frequency of scores 3 and 12. This pattern was similar when examining the distribution of histology grades in colons from *Study 1*, where 42% were free of microscopic lesions (Fig. 7B). In contrast, only 24% of small intestinal sections received a histology grade of 0. Thus, more than half of small intestines with a normal macroscopic appearance may still exhibit some signs of pathology upon histological assessment.

**Figure 7.**
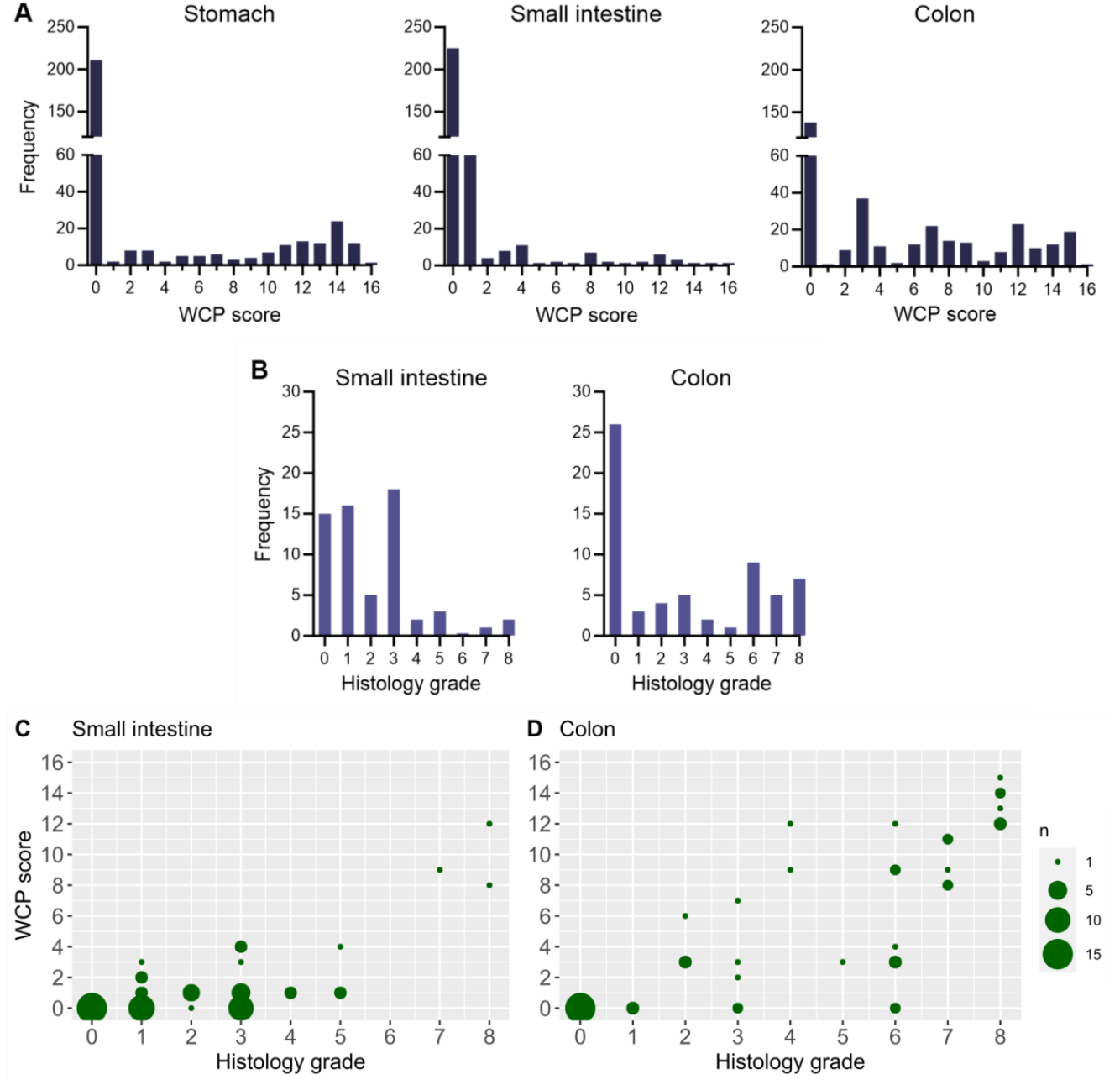
Weighted and cumulative point (WCP) score and histopathological grade observations. A) Frequency of macroscopic WCP scores by organ (*n* = 333). B) Frequency of histopathological grades by organ (*n* = 62). C) Correlation between the macroscopic WCP scoring of the whole organ and the histopathologic assessment of the worst lesion site (*n* = 62).

To investigate this further, we correlated individual macroscopic WCP scores with histology grades from *Study 1* (Fig. 7C). Notably, 19 small intestines (31% of the subset) were observed normal or hyperemic at the macroscopic inspection (WCP score 0-1) but received a histology grade 3-5, ranging from local lifting or disruption of villus tip epithelium to villus stunting. Additionally, seven colons (11% of the subset) showed local or pronounced mucosal ulceration at the histopathological assessment (grade 5-6) but exhibited no macroscopic visible necrosis. Collectively, the histological grading of intestinal lesions can complement the pathological assessment, revealing intestinal damage that is not visible through gross inspection. The extended range of the WCP score demonstrated a better visually apparent correlation with histopathology when compared to the categorical 6-grade score, particularly in colons (Supplementary Fig. S3).

### NEC sensitivity is affected by birth weight

The median birth weight across all studies was 960 grams ± 255 grams. When correlating the individual WCP scores to birth weights, we saw a slight rise in lesion levels as the weight increased (Supplementary Fig. S4). This was statistically evident in the small intestine and colon, but not in stomachs (*P-*values of 0.02, 0.05, and 0.3, respectively). The distribution of sexes across studies was equal (48% females) and, as previously reported, the sex had no statistical impact on the pathology levels in any gastrointestinal region (Supplementary Fig. S5) [14]. This indicates that larger preterm piglets are slightly more sensitive to NEC than smaller littermates in the included studies. Moreover, in the examination of WCP scores, it may be beneficial to include birth weight in the statistical model to enhance statistical power.

### The weighted and cumulative point system distinguishes two NEC-inducing diets

To illustrate the application of the WCP scoring system, we examined the NEC-inducing effect of a gently processed (GP) home-mixed formula in comparison to a harshly processed (HP) ready-to-drink formula. Employing a proportional ordered logistic regression model with formula, birth weight, and study origin as covariates, we found that the HP formula induced higher macroscopic scores, particularly in colons (Fig. 8A). Here, GP feeding resulted in a median score of 3 points, while HP feeding increased the median score to 7 points. Due to the low level of small intestinal lesions, both formula groups had a median score of 0 in this region. Nevertheless, a statistical increase was observed, as 8% of GP milk-fed pigs and 21% of HP milk-fed pigs received a score of 4 or above. The formula did not affect the level of stomach lesions. The same patterns were observed when analyzing the categorical 6-grade scores (Supplementary Fig. S6). When exploring specific pathologies, the HP formula led to a higher incidence of pneumatosis intestinalis in the small intestines (Fig. 8B). Hemorrhage and necrosis incidences also tended to be elevated. Similarly, but with greater significance, incidences of these three pathologies were notably higher in colons from the HP milk-fed group. When comparing lesion cytokine levels, only IL-1β in colons showed a significant increase when fed the HP formula (Fig. 8C). In conclusion, the harshly processed milk formula induced a higher level of NEC-like lesions compared to the gently processed formula. This included a higher level of pneumatosis intestinalis and necrosis, both of which are hallmarks of NEC.

**Figure 8.**
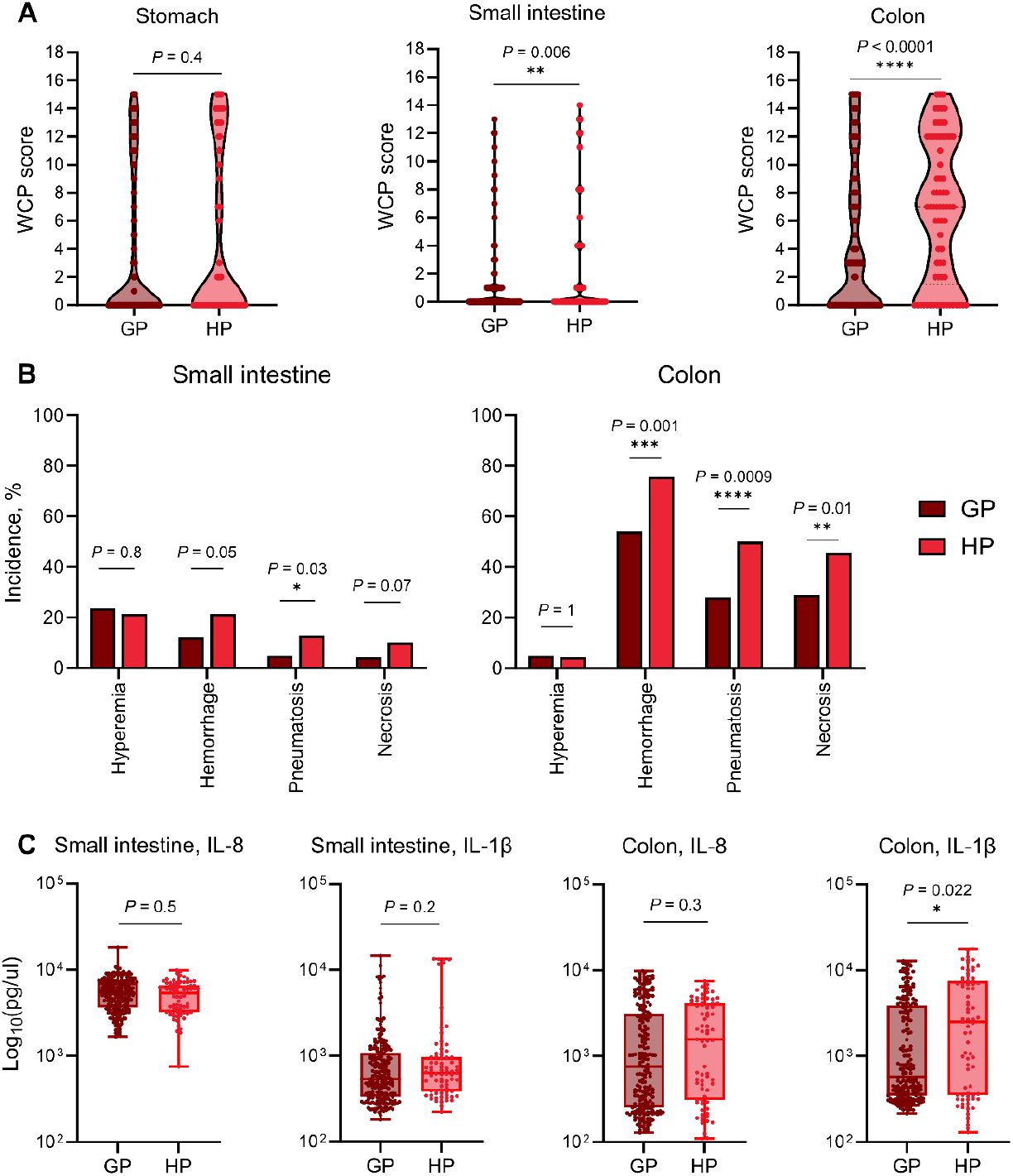
Comparison of a gently processed (GP, *n* = 263) and harshly processed (HP, *n* = 70) milk formula on gut inflammation. A) Cumulative and weighted point (WCP) scores of each gastrointestinal segment. B) Incidence of each pathology type in GP milk-fed and HP milk-fed piglets. C) Cytokine measurements in local lesions from small intestines and colons. IL = interleukin.

## Discussion

The piglet NEC model is based on preterm birth and formula feeding, leading to the spontaneous development of intestinal lesions without the introduction of artificial physiological stressors. The overall pathophysiology thus resembles that of NEC and the lesions exhibit the same gross - and histopathological hallmarks. Therefore, the piglet has the closest resemblance to human NEC in terms of constructive - and face validity, compared to rodent models [8–10]. Irrespectively, the use of an NEC animal model allows for a systematic and controlled assessment of treatment effects directly on the level of intestinal lesions. The accuracy of such an assessment is almost as important as the model’s validity to ensure proper interpretation of the experiment.

The systematic necropsy of all animals allows careful inspection of pathology across the GI tract. Here, the categorical 6-grade NEC scoring may be overly simplistic in accurately capturing the diversity of these pathologies. Grades 3-4 are defined by hemorrhage and grades 5-6 by necrosis and/or pneumatosis intestinalis. As a result, this system fails to describe lesions, where all three pathology types are present simultaneously and thereby fails to discriminate these severe cases from lesions with, for instance, only a few pneumatoses. These considerations led to the development of the WCP scoring system where pathologies are reported separately and subsequently weighted and combined in one cumulative NEC score. This features an expanded score range, particularly addressing the heterogeneity of severe lesions, and the possibility to explore each pathology type further.

NEC diagnosis is associated with elevated levels of various proinflammatory cytokines within the intestinal tissues [6, 15]. Particularly, NEC tissue specimens from preterm infants and piglets have shown a rise in IL-1β and IL-8 [16, 17]. The increasing WCP score correlated well with elevated levels of these two pro-inflammatory cytokines. This indicates that a higher WCP score reflects a higher degree of lesion-associated inflammation. Only small intestinal IL-8 did not correlate to higher pathology levels, which may be due to the constitutive expression of this cytokine in immature small intestinal enterocytes [18]. The absence of a correlation between hyperemia and elevated cytokine levels confirms the limited association between hyperemia and intestinal inflammation. This supports the practice of assigning only a marginal weight to hyperemia, irrespective of its extent. Although only analyzed in a subset of samples, the histopathological grade system further supported the macroscopic findings. As with cytokine measurements, this was assessed in lesion biopsies and thus does not reflect the overall level of pathology. As such, a high histopathological grade may exacerbate the severity compared to the WCP score by not evaluating the whole GI segment. However, several organs with a normal macroscopic appearance were found affected upon histopathological assessment. Consequently, the histologic assessment not only validates and supports the NEC score but also serves to identify discrete mucosal lesions that are not easily recognized macroscopically.

Although the validity of the preterm piglet NEC model is superior, and the model is recognized as being the most clinically relevant [8–10], some discrepancies do exist. During the past years, we have observed that piglets born with a higher birth weight tend to develop more NEC. To account for this, we fed all pigs with daily volumes adjusted to metabolic rates (mL/kg^0.75^), whereby small pigs received relatively larger volumes. Still, the current dataset suggests a subtle yet statistically significant correlation between higher birth weight and an elevated WCP score. This aspect contradicts findings in preterm infants, where low weight is a clear risk factor for NEC [19]. This may reflect an interspecies difference in NEC susceptibility. Alternatively, as postulated by Carter et al. [20], it can indicate that comorbidities (e.g. sepsis and mechanical ventilation) play a more significant role than birth weight alone. Furthermore, NEC-like lesions in preterm piglets manifest primarily in the colon and to a lesser extent in the stomach, with infrequent lesions in the small intestine. In preterm infants, stomach lesions are rarely observed, whereas ileum and proximal colon are most frequently affected [21]. Interestingly, we do not see a difference between the two infant formulas in terms of stomach NEC score, which is the case for the small intestine and colon. This could point to different lesion etiologies, which should be considered when interpreting lesion manifestation across the GI tract in response to experimental treatments.

The comparison of milk formulas mainly functions to demonstrate the utility of the WCP score in comparing NEC lesions between groups and investigating differences in separate pathologies related to NEC. The formulas differed in terms of the major carbohydrate source (maltodextrin vs. lactose) and processing (pasteurization and spray-drying vs. ultra-high temperature treatment). Although the gently processed formula contained more maltodextrin, which is documented to be much more NEC-promoting than lactose in preterm piglets [22], the harshly processed formula induced a higher WCP score, characterized by increased occurrence of hemorrhage, pneumatosis intestinalis, and necrosis.

## Conclusion

We have developed and validated a new weighted and cumulative scoring system to access gastrointestinal pathology in the preterm piglet model of NEC. The use of this scoring system may improve the accuracy of pathological assessment and increase the efficiency of preclinical NEC experiments.

