## Supplementary materials for "A weighted and cumulative point system for accurate scoring of intestinal pathology in a piglet model of necrotizing enterocolitis"

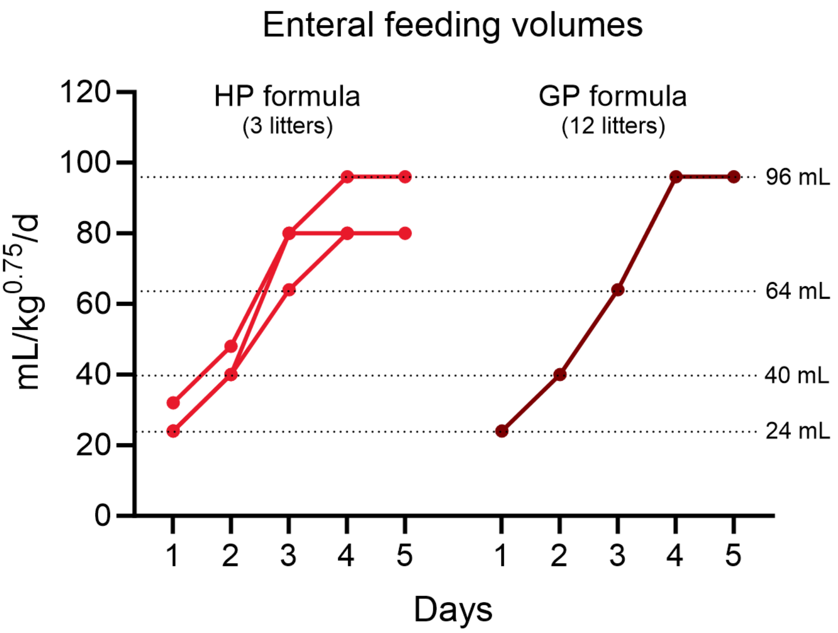


**Supplementary** Figure S1: Overview of daily enteral feeding volumes between piglet litters fed a harshly processed (HP) or a gently processed (GP) milk formula.

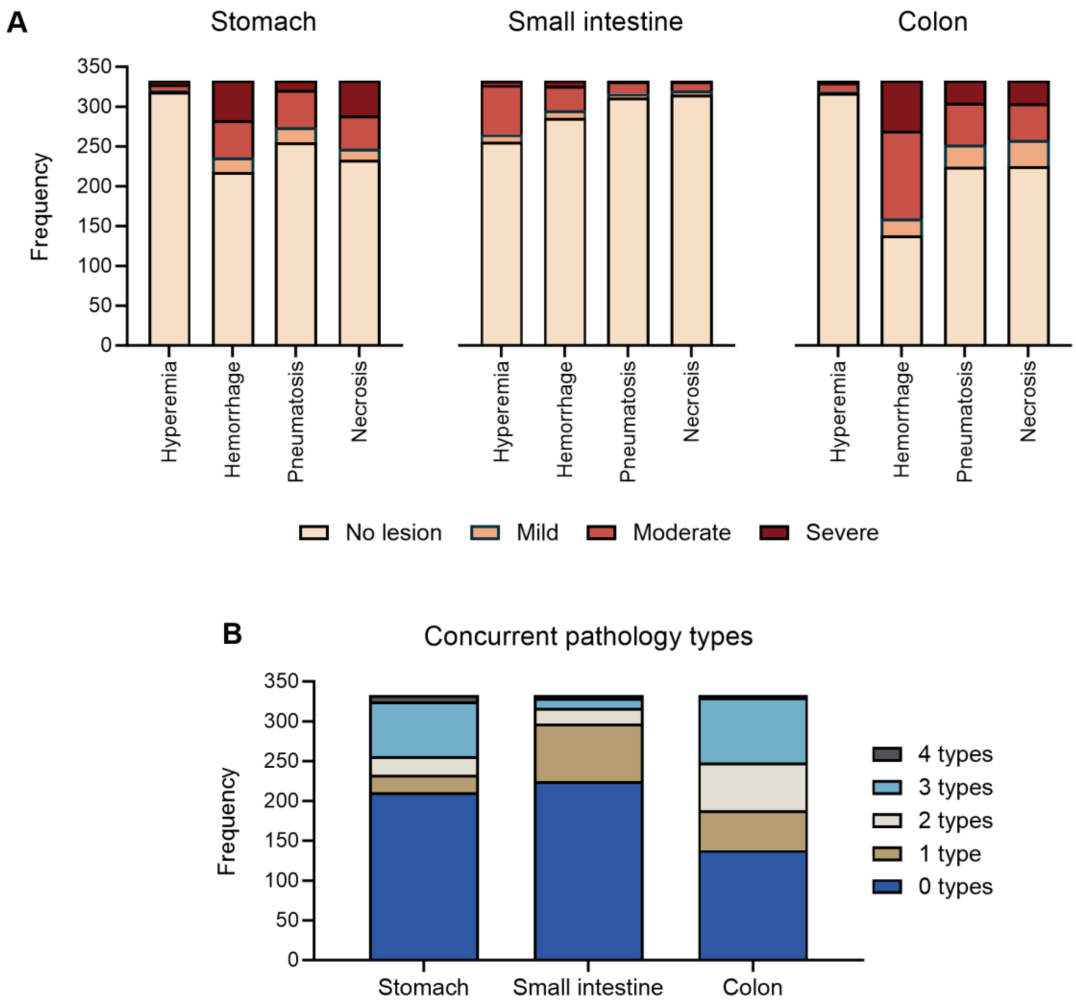


**Supplementary** Figure S2: Overview of the four pathology types; hyperemia, hemorrhage, pneumatosis intestinalis, and necrosis (*n* = 333). A) Distribution of each type and extent in the stomach, small intestine, and colon. B) Overview of the number of concurrent pathology types in each gastrointestinal segment.

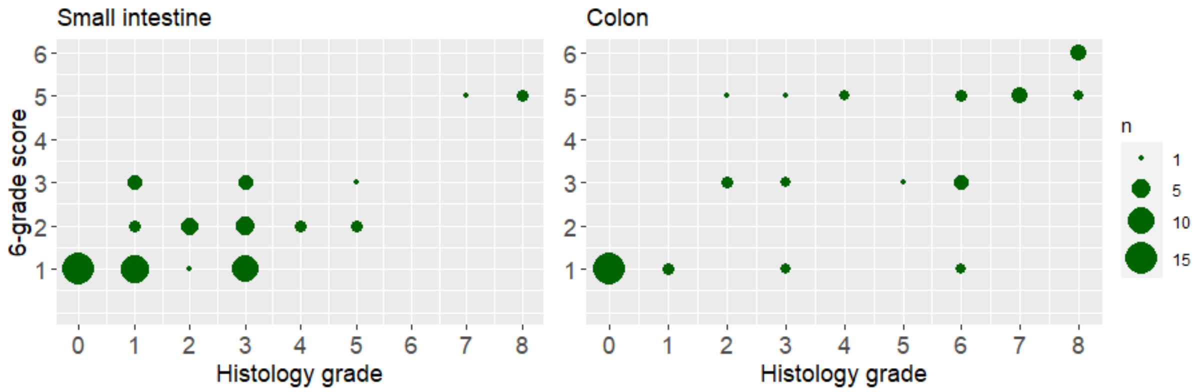


**Supplementary** Figure S3: Correlation between the macroscopic 6-grade score of the whole organ and the histopathologic assessment of the worst lesion site (*n* = 62).

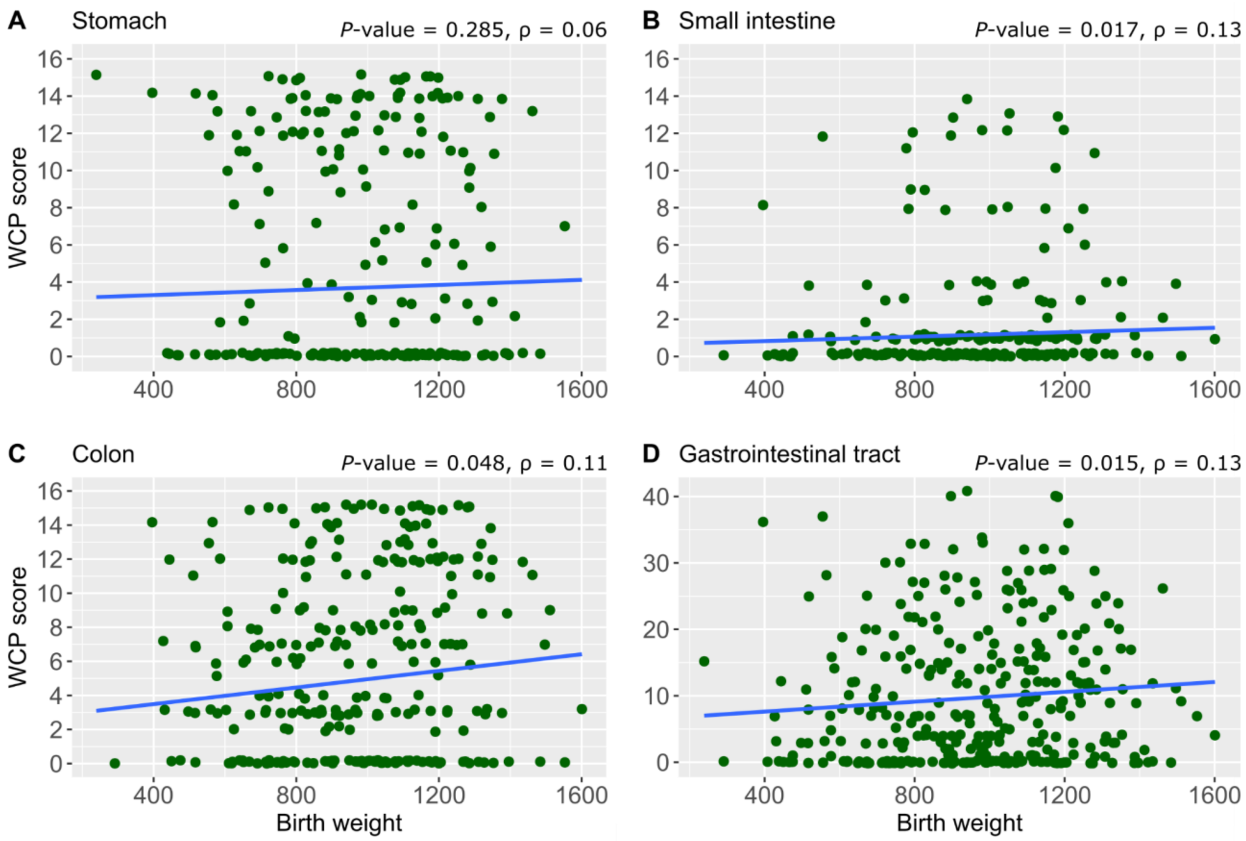


**Supplementary** Figure S4: Correlation between weighted and cumulative point (WCP) scores and birth weight using Spearman's rho (ρ) in gastrointestinal sections separately; the stomach (A), small intestine (B), and colon (C), as well as the sum across the gastrointestinal tract (D). *n* = 333 pigs.

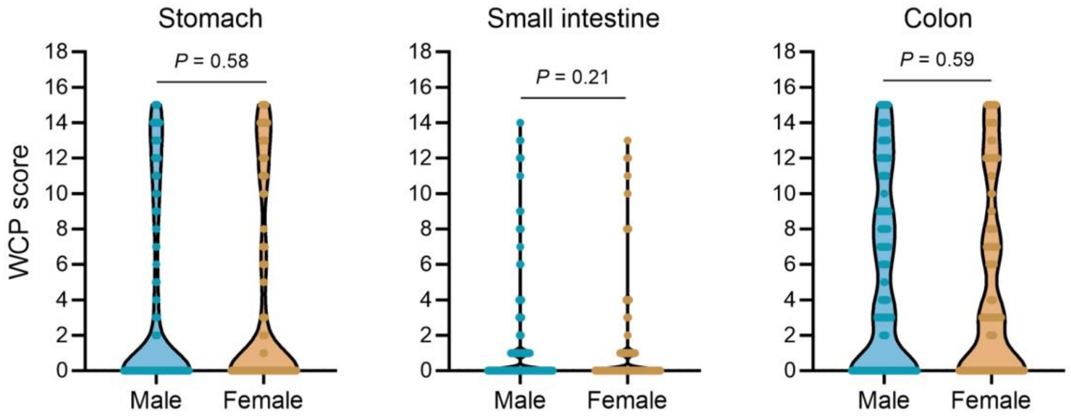


**Supplementary** Figure S5: Comparison of cumulative and weighted point (WCP) scores between males and females in stomachs, small intestines, and colons ($n$ = 333).

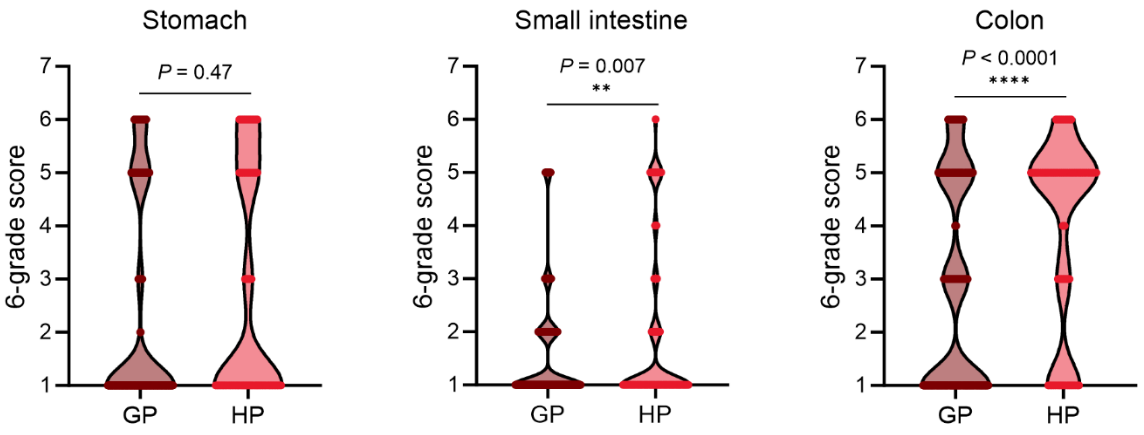


**Supplementary** Figure S6: Comparison of a gently processed (GP, $n$ = 263) and harshly processed (HP, $n$ = 70) milk formula on gut inflammation using the categorical 6-grade scoring system.

**Supplementary** Table S1: Formula energy compositions.

| | GP formula | HP formula |
|---|---|---|
| Energy (kJ/L) | 3661 | 3922 |
| Protein (g/L) | 66 | 46 |
| - Whey protein (g/L) | 53 | 46 |
| Carbohydrate (g/L) | 47 | 78 |
| - Lactose (g/L) | 5 | 55 |
| - Maltodextrin (g/L) | 42 | 23 |
| Fat (g/L) | 48 | 49 |
| - Saturated fats (g/L) | 36 | 25 |
| - Monounsaturated fats (g/L) | 6 | 11 |
| - Polyunsaturated fats (g/L) | 3 | 8 |

GP = gently processed, home-mixed formula. HP = harshly processed ready-to-drink formula supplemented with extra fat and protein.

**Supplementary** Table S2: Cytokine levels affected by hyperemia, hemorrhage, pneumatosis intestinalis, and necrosis.

| | **IL-8** | | **IL-1β** | |
|---|---|---|---|---|
| | **Estimate**[a] (95% CI) | ***P*-value** | **Estimate**[a] (95% CI) | ***P*-value** |
| **Small intestine** | | | | |
| Hyperemia | 0.71 (0.63, 0.80) | 7.49e-08 **** | 1.09 (0.86, 1.39) | 0.475 |
| Hemorrhage | 0.94 (0.78, 1.13) | 0.487 | **1.67** (1.15, 2.42) | 0.007 ** |
| Pneumatosis | 0.61 (0.43, 0.86) | 0.005 ** | 0.81 (0.41, 1.60) | 0.542 |
| Necrosis | **1.92** (1.36, 2.72) | 0.0003 *** | **4.62** (2.32, 9.18) | 2.02e-05 **** |
| **Colon** | | | | |
| Hyperemia | 0.97 (0.58, 1.30) | 0.919 | 0.81 (0.50, 1.32) | 0.3996 |
| Hemorrhage | **2.41** (1.81, 3.22) | 5.22e-09 **** | **2.59** (1.97, 3.40) | 5.40e-11**** |
| Pneumatosis | **2.04** (1.54, 2.70) | 1.22e-06 **** | **2.21** (1.69, 2.89) | 1.74e-08**** |
| Necrosis | **2.66** (2.00, 3.53) | 6.43e-11 **** | **2.41** (1.84, 3.16) | 5.16e-10**** |

[a] The estimate shows the relative difference in cytokine levels when the pathology type is present. For instance, an estimate of 2 indicates that the cytokine level doubles in the presence of the pathology. Pathology types that significantly *increase* cytokine levels are highlighted in bold. CI = confidence interval.
